# Architecture Matters: Design Rules for Multigene IDO1/PD-L1 Cassettes in Human Skin Cells

**DOI:** 10.64898/2026.03.28.714644

**Authors:** Hamid Reza Karbalaei-Heidari, Rahele Daraeinejadfard, Afshin Raouf, Sarvesh Logsetty, Rae Spiwak, Song Liu, Nediljko Budisa

## Abstract

Allogeneic cell therapies require the coordinated expression of multiple immunomodulatory genes, yet multigene circuits that function in permissive cell lines often fail in differentiated human tissues for unclear reasons. Here, we systematically dissect how transcriptional architecture governs functional immunoregulation in engineered human keratinocyte and fibroblast lines. Using site-specific large-cargo integration (eePASSIGE) as an enabling tool, we determined that genomic insertion efficiency was not the limiting factor for phenotype; rather, promoter arrangement and gene order dictated expression hierarchy. A single-promoter EF1α-IDO1-T2A-GFP design that expressed robustly in HEK293T cells was nearly silent in skin-derived cells, preventing reporter-based enrichment. In dual- and tri-modular cassettes, we observed severe transcriptional interference: a downstream CMV promoter driving GFP or PD-L1/iCasp9 (via EMCV-IRES) markedly suppressed the upstream EF1α-IDO1 unit, despite intact integration (resulting in ∼175–625-fold attenuation), demonstrating strong promoter interference within the circuit. Functionally, co-culture assays revealed a hierarchical immunomodulatory logic: high IDO1 expression proved to be a requisite threshold for T-cell suppression, whereas PD-L1 provided measurable benefit only against highly activated, PD-1^+^ T cells in vitro. Collectively, these data establish a site-specific framework for generating immune-tuned skin cells and define essential design rules for avoiding promoter interference in next-generation translational skin substitutes.

## 1. Introduction

Severe burns and chronic wounds create an urgent demand for readily available, clinically practical skin substitutes that can be deployed rapidly and at scale [1, 2]. Although autologous split-thickness skin grafting remains the standard of care, it is fundamentally constrained by donor-site morbidity and limited tissue availability, particularly in patients with extensive total body surface area burns [3]. Allogeneic skin constructs and other “off-the-shelf” approaches offer a solution to supply limitations; however, their durable application is restricted by host immune rejection, which compromises engraftment, persistence, and long-term function [4]. A key translational opportunity, therefore, is to engineer graft-forming cells that can autonomously attenuate alloreactive immune responses locally, reducing reliance on systemic immunosuppression and improving durability in clinically relevant settings [5].

The development of a universal skin substitute requires the coordinated engagement of multiple immunomodulatory mechanisms that mimic endogenous pathways of peripheral tolerance. One compelling axis is Indoleamine 2,3-dioxygenase 1 (IDO1), an enzyme that catalyzes the rate-limiting step of tryptophan catabolism into kynurenines. Local tryptophan depletion and kynurenine signaling suppress T-cell expansion through amino-acid starvation responses (e.g., GCN2 signaling/activation) and promote regulatory phenotypes in antigen-presenting and lymphoid compartments [6-8]. In principle, embedding IDO1 activity within a graft microenvironment offers a metabolic checkpoint that dampens effector responses without requiring systemic drug exposure. Concurrently, Programmed Death-Ligand 1 (PD-L1; CD274) provides an orthogonal and clinically validated checkpoint pathway. The engagement of PD-L1 with PD-1 on activated T cells inhibits proximal T-cell receptor signaling and downstream growth programs, limiting effector function and promoting exhaustion under persistent stimulation [9, 10]. Mechanistically distinct, IDO1 (metabolic checkpoint) and PD-L1 (inhibitory checkpoint) are poised to act additively within engineered skin constructs, particularly under inflammatory conditions in which alloreactivity is most potent.

However, achieving clinically meaningful immune modulation in engineered cells requires stable, predictable, and safe gene expression. Site-specific integration into genomic “safe-harbor” loci, such as AAVS1, is widely used to mitigate position effects and reduce the risk of insertional mutagenesis relative to random integration [11]. Recently, large-cargo strategies coupling prime editing with serine integrases have enabled the installation of landing pads and the subsequent recombination of multi-kilobase donor payloads. Unlike traditional homology-directed repair, these approaches - exemplified by PASTE and evolved variants such as eePASSIGE - avoid the generation of double-strand breaks that drive indels and chromosomal rearrangements [12, 13]. While these platforms provide a practical route to integrate multigene payloads into defined loci, achieving therapeutic thresholds remains challenging, as transgene stability is highly sensitive to cell-type expression constraints and the specific configuration of promoter architecture.

Beyond integration strategies, translational designs necessitate control layers that address safety and metabolic fitness. An inducible apoptosis module, such as iCasp9, functions as a drug-activated safety switch, enabling the rapid ablation of engineered cells upon dosing with a bioinert small-molecule dimerizer like rimiducid (AP1903) [14, 15]. Furthermore, because constitutive IDO1 expression can impose a metabolic fitness cost, doxycycline-inducible (Tet-On) regulation provides a mechanism to restrict expression to experimental windows or peri-engraftment periods [16, 17]. Crucially, multigene systems introduce an underappreciated layer of complexity: cassette architecture itself - encompassing promoter choice, strength balance, orientation, and insulation - can drive transcriptional interference or silencing, ultimately determining whether engineered cells meet functional targets.

In this study, we engineered human keratinocytes (HaCaT) and fibroblasts (HDFn-hTERT) to stably express IDO1 and PD-L1, coupled to inducible control and an iCasp9 safety switch, using a modular site-specific integration workflow. We first benchmarked prime editing-directed integrase configurations for large-cargo insertion in HEK293T cells, identifying eePASSIGE as the most efficient platform. We then evaluated how cassette architecture shapes expression stability and functional output across cell types, quantified transgene expression and safety-switch performance, and validated immunomodulatory activity in T-cell co-culture assays. Together, this work establishes design principles for multigene immune-modulating cassettes in skin-forming cells and provides a modular framework to accelerate the development of next-generation translational skin substitutes.

## 2. Results

### 2.1. Benchmarking large-fragment insertion identifies eePASSIGE as the most efficient platform in HEK293T cells

To identify the optimal strategy for large-cargo integration, we benchmarked three prime editing-directed integrase platforms - PASTE, eePASTE, and eePASSIGE - at a shared genomic locus in HEK293T cells. To ensure consistent delivery and minimize variation associated with electroporation, all constructs were introduced via lipid-mediated transfection using Lipofectamine 3000. Integration efficiency was quantified as the fraction of correctly modified cells using reporter assays and locus-spanning PCR, with site-specific junctions confirmed by Sanger sequencing. Across independent biological replicates (n =3), eePASSIGE yielded the highest targeted integration rates (median 5%) compared to PASTE (2%) and eePASTE (3%). Cellular viability remained comparable across all transfection conditions. Based on this superior editing performance, we selected eePASSIGE as the platform for all downstream engineering in skin-derived cell lines.

### 2.2. Ribosomal skipping inefficiency and promoter interference limit cassette performance in skin-derived cells

To establish stable immunomodulatory lines, we first evaluated a single transcription unit (TU) cassette (Construct 637: EF1*α* → IDO1-T2A-GFP). While this design yielded robust reporter expression and successful sorting in HEK293T cells, it failed to produce detectable GFP signals in HaCaT and HDFn lines following nucleofection. Given that the EF1*α* promoter is generally active in these lineages, the absence of fluorescence suggests a failure of the T2A ribosomal skipping sequence to function efficiently in these specific cell types. Inefficient skipping results in the translation of a steric-hindered IDO1-GFP fusion product rather than discrete proteins, leading to misfolding and rapid proteasomal degradation of the reporter [18].

To overcome this translational bottleneck and decouple reporter detection from therapeutic gene expression, we transitioned to a dual-promoter design (Construct 715: EF1 *α* → IDO1-BGH; CMV → GFP-SV40pA). This strategy successfully restored GFP expression, enabling the isolation of clonal populations via single-cell sorting. However, quantification of transgene levels in these expanded clones revealed a critical decoupling effect. RT-qPCR analysis, normalized to the geometric mean of *GAPDH* and *B2M*, showed that while 637-HEK clones maintained high IDO1 expression (7% of reference transcripts), the sorted 715-HaCaT clones exhibited markedly suppressed IDO1 levels (0.04% of reference; ∼175-fold reduction) (Fig. 1). This discordance indicates that while the separate CMV unit drove sufficient GFP for sorting, the upstream EF1*α*-IDO1 cassette was subject to severe transcriptional interference (promoter occlusion) by the potent downstream CMV module in this uninsulated tandem arrangement [19, 20].

**Fig. 1.**
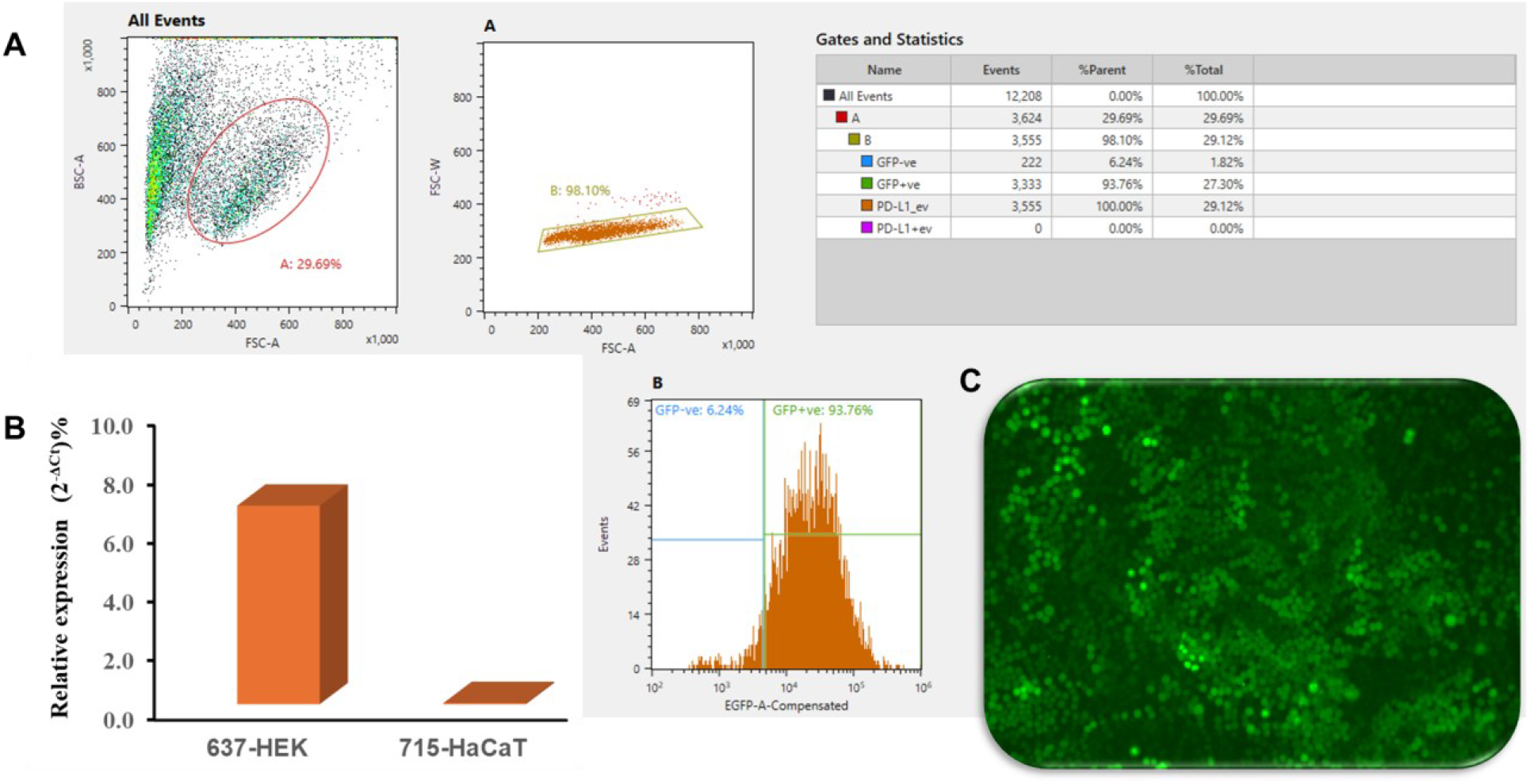
Vector architecture determines the correlation between reporter intensity and therapeutic transcript levels. **(A)** Representative flow cytometry histogram showing GFP fluorescence intensity in the 715-HaCaT cell population, confirming widespread and robust transgene expression driven by the downstream CMV promoter across the bulk population. **(B)** Quantitative real-time PCR (qPCR) analysis of IDO1 mRNA expression in 637-HEK293T and 715-HaCaT clonal cells, presented as a percentage of the geometric mean of two reference genes (*GAPDH* and *B2M*) (ref. geo %) to enable normalized cross-cell-line comparison. 637-HEK293T cells, which carry an EF-1α-IDO1-T2A-GFP polycistronic cassette, exhibited an IDO1 transcript level of 6.75% relative to the reference gene geometric mean. In contrast, 715-HaCaT cells, which harbor a bicistronic dual-promoter cassette (EF-1α-IDO1-CMV-GFP), showed a dramatically reduced IDO1 mRNA level of 0.04% - representing an approximately 170-fold reduction - despite confirmed stable genomic integration of the full-length construct. **(C)** Representative fluorescence microscopy images of GFP expression in 715-HaCaT cells. Notably, 715-HaCaT cells displayed GFP fluorescence intensity that was qualitatively equal to or brighter than that observed in 637-HEK293T cells, despite the latter relying on T2A-mediated ribosomal skipping from a single EF-1α-driven transcript. The strong GFP signal in 715-HaCaT cells confirms that the integrated locus is transcriptionally accessible and that the CMV promoter is highly active in these keratinocytes. Critically, however, the upstream EF-1α promoter driving IDO1 within the same construct is nearly silent, yielding transcript levels ∼170-fold lower than those observed in HEK293T cells. This discordance between two promoters residing within the same integrated cassette implicates promoter interference as a primary mechanism. Regardless of mechanism, this finding demonstrates that dual-promoter cassette architecture does not guarantee coordinate expression of both transgenes and that promoter–promoter interactions within multigene constructs can create silent bottlenecks in which one therapeutic gene is genomically present and flanked by an active expression unit yet remains functionally silent in differentiated human cell types.

### 2.3. CMV-driven EMCV-IRES architecture provides efficient, balanced expression in primary-like cell lines

To establish a baseline for checkpoint inhibitor expression, we evaluated Construct 702 (EF1*α* → PD-L1–IRES–GFP). In contrast to the ribosomal skipping inefficiency observed with T2A peptides in these lineages, the EMCV IRES system successfully mediated the internal ribosome entry required for bicistronic translation. This architecture generated a single transcript that was efficiently translated into both PD-L1 and GFP, resulting in comparable high-level expression of both the therapeutic cargo and the reporter. In HEK293T cells, this configuration drove potent PD-L1 transcript levels (∼486% relative to reference genes).

We then assessed a dual-promoter IDO1/PD-L1 cassette (Construct 706: EF1*α* → IDO1-BGH; CMV → PD-L1-SV40pA) to determine if independent regulation could be achieved. However, quantitative PCR analysis revealed a severe transcriptional imbalance. While PD-L1 expression remained remarkably high (∼339% of reference) driven by the CMV promoter, the upstream EF1*α*-driven IDO1 levels were drastically reduced to ∼0.4% (Fig. 2). Crucially, this pattern of robust PD-L1 expression and suppressed IDO1 was conserved across both HEK293T and HaCaT cell lines, resulting in a ∼625-fold transcript imbalance (PD-L1:IDO1). Together with the reporter data, these findings indicate that: (i) PD-L1 is readily expressed under CMV control using EMCV-IRES architectures in both kidney and skin lineages, whereas (ii) the upstream IDO1 locus is highly sensitive to vector architecture, likely undergoing transcriptional silencing (promoter occlusion) induced by the adjacent, highly active CMV promoter [21] (Scheme 1).

**Fig. 2.**
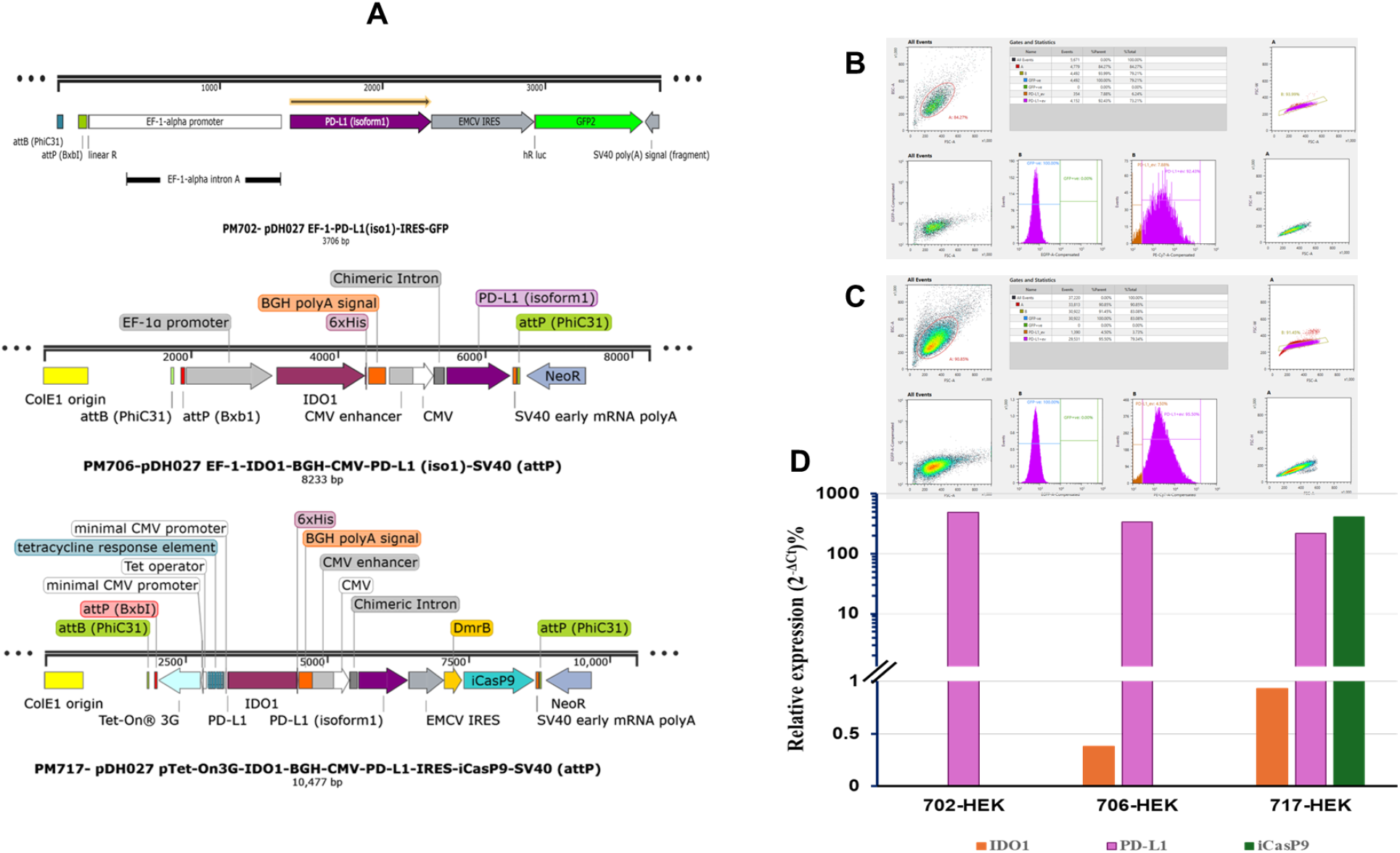
Construction and validation of multi-cistronic vectors for co-expression of immunomodulatory and safety-switch genes. **(A)** Schematic linear maps of the engineered constructs generated using SnapGene. The panel details the vector architectures: **702** (Control: EF-1*α*-IRES-GFP), **706** (Dual-Promoter: EF-1*α*-IDO1 / CMV-PD-L1), and **717** (Tri-genic: EF-1*α*-IDO1 / CMV-PD-L1-IRES-iCasP9). Note the use of distinct promoters (EF-1*α* and CMV) to separate *IDO1* and *PD-L1* expression, and the IRES element in 717 to couple *iCasP9* translation to the *PD-L1* transcript. **(B-C)** Flow cytometric validation of surface PD-L1 protein expression in stably transfected HEK293T cells using anti-CD274-PE-Cy7 antibody. Panel **(B)** displays the expression profile of the 706-HEK population, while **(C)** displays the 717-HEK population. **(D)** Quantitative real-time PCR (qPCR) analysis of relative mRNA expression levels of IDO1, PD-L1, and iCasp9 in 702-HEK293T, 706-HEK293T and 717-HEK293T cells, presented as a percentage of the geometric mean of two internal reference genes (*GAPDH* and *B2M*) (ref. geo %).

**Scheme 1:**
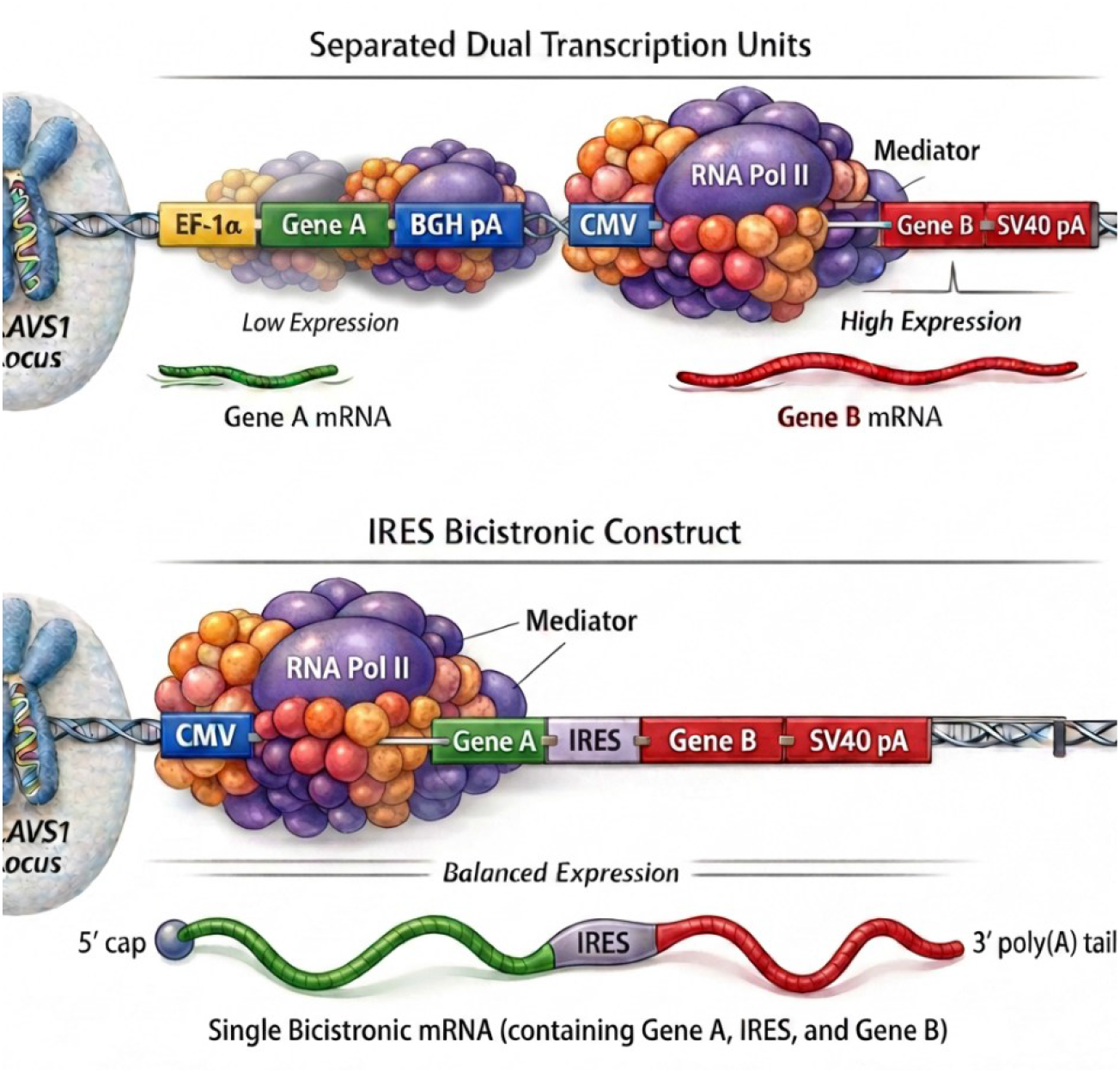
Impact of vector design on transcriptional consistency. (Top) Separated dual transcription units results in promoter occlusion, a phenomenon where transcriptional interference between adjacent promoters disrupts the stoichiometric balance of gene expression. (Bottom) In contrast, A bicistronic design utilizing an IRES-containing single transcription unit ensures coupled expression, maintaining a consistent level of transcription for both genes of interest.

### 2.4. Tri-modular cassette drives robust PD-L1 and iCasp9 expression but fails to trigger apoptosis in HEK293T cells

To integrate both immunomodulation and a safety switch, we evaluated a tri-modular design (Construct 717: EF1 *α* → IDO1-BGH; CMV → PD-L1–IRES–DmrB-iCasp9–SV40pA). In HEK293T cells, RT-qPCR analysis of the integrated gene cassette in AAVS1 locus revealed a familiar pattern of transcriptional interference: upstream IDO1 expression was severely attenuated (∼1% of reference transcripts), while the downstream CMV-driven unit produced high levels of both PD-L1 (∼220%) and iCasp9 (∼410%).

Despite abundant *iCasp9* mRNA, treatment with the chemical inducer of dimerization, rimiducid (in the range of 1 nM–1 μM), failed to trigger apoptosis above baseline levels. This functional disconnect likely stems from two synergistic factors. First, our construct utilized the full-length Caspase-9 sequence, retaining the N-terminal CARD domain. This domain can sterically hinder the efficient drug-induced dimerization required for activation, a limitation overcome in later-generation “truncated” iCasp9 variants [15, 22]. Second, HEK293T cells express adenoviral E1B proteins (specifically E1B-19K), which function as potent Bcl-2 homologs to buffer against mitochondrial apoptosis, effectively raising the threshold required for Caspase-9/3-mediated cell death [23, 24].

### 2.5. IDO1 expression drives robust T-cell suppression, whereas PD-L1 efficacy is contingent on T-cell activation state

To validate the functional impact of our engineered immunomodulatory cassettes, we performed co-culture assays with human PBMCs. Notably, construct 637-integrated HEK cells, which maintain robust IDO1 expression, demonstrated significant suppression of T-cell proliferation compared to wild-type controls, confirming the potency of the metabolic checkpoint in this system (Fig.3).

**Figure 3.**
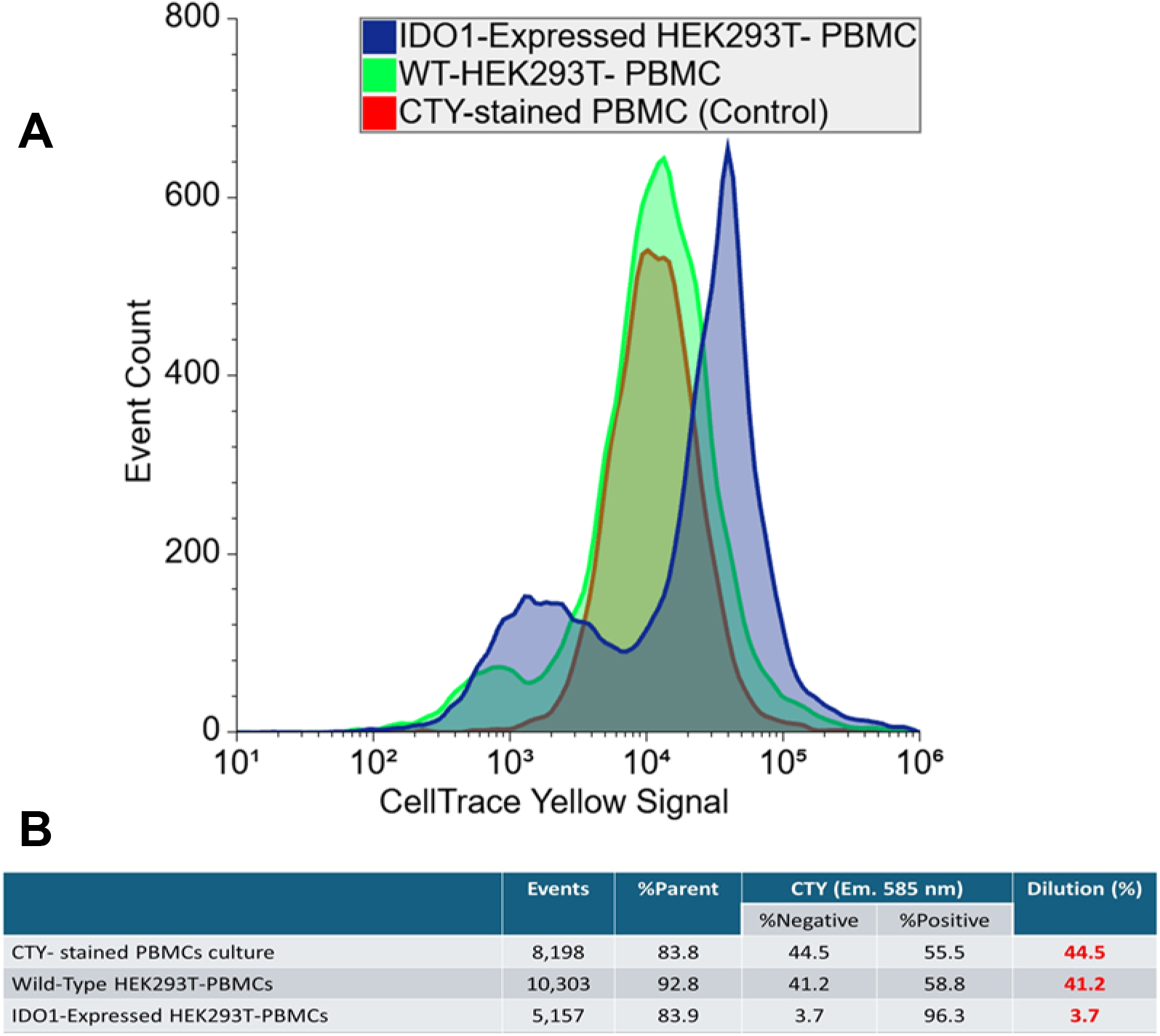
IDO1-expressing HEK293T cells suppress T-cell proliferation in a mixed lymphocyte co-culture assay. Peripheral blood mononuclear cells (PBMCs) were labeled with CellTrace™ Yellow (CTY) proliferation dye and cultured for 72 hours under three conditions: PBMCs alone (control), PBMCs co-cultured with wild-type HEK293T cells (WT-HEK293T), and PBMCs co-cultured with IDO1-expressing HEK293T cells (clone 637-HEK293T). T-cell proliferation was assessed by flow cytometric analysis of CTY dye dilution as a readout of successive cell divisions. Flow cytometry gating strategy identified lymphocytes by forward scatter (FSC) versus side scatter (SSC), followed by the selection of CD3^+^ T cells (anti-CD3-PerCP-Cy5.5) and CD8^+^ cytotoxic T cells (anti-CD8-PE-Cy7). A) The accompanying histogram overlay plots Total Events versus CTY fluorescence signal across all three culture conditions. Peaks shifted to the right (CTY-high) represent the undivided parental population, while peaks shifted to the left (CTY-low) indicate proliferating cells that have undergone dye dilution. **(B)** Quantification of T-cell proliferation. Co-culture with 637-HEK293T cells resulted in marked immune suppression (∼ 4.0% dilution) consistent with IDO1-mediated tryptophan catabolism. The percentage of proliferating (diluted) cells for each condition is detailed in the table.

In contrast, construct 702-HEK cells (high PD-L1) did not exhibit enhanced suppression relative to controls under standard activation conditions. This likely reflects the specific activation threshold required for checkpoint inhibition: effective PD-L1 signaling necessitates the substantial upregulation of its receptor, PD-1, on activated T-cells. In our assays, donor variability— specifically high baseline Regulatory T-cell (T-reg) populations in control samples—or insufficient T-cell activation may have masked the suppressive potential of the PD-L1 axis. Consequently, demonstrating the specific efficacy of PD-L1 requires stimulation conditions optimized to induce high-level PD-1 expression on effector T-cells, distinct from the metabolic starvation mechanism of IDO1.

Furthermore, our results highlight the critical influence of cytokine context on suppression assays. We observed that high-dose exogenous IL-2 (e.g., 1000 U/mL) abrogated the IDO1-mediated suppression in 637-HEK co-cultures, consistent with strong IL-2R-driven STAT5 and PI3K–Akt– mTORC1 signaling overriding amino acid–starvation checkpoints [7, 10].

However, these *in vitro* constraints underscore the necessity of testing in a physiological setting. The inflammatory microenvironment of severe burns and chronic ulcers is naturally characterized by a high frequency of activated effector T cells that significantly upregulate PD-1. In this clinical context, we anticipate that our engineered constructs—particularly those expressing PD-L1—will demonstrate superior potency compared to static co-cultures, as the local expression of PD-L1 will directly engage these naturally activated T-cell phenotypes.

## 3. Discussion

The engineering of universally transplantable skin substitutes is commonly approached as a problem of efficient genome integration and sufficient transgene expression. Our results indicate that this framing is incomplete. In this study, we established a modular framework for engineering hypoimmunogenic skin cells, identifying eePASSIGE as a superior platform for large-cargo integration. However, our central finding is that successful genomic insertion is merely the first step; functional outcome was determined not by editing strategy but by transcriptional architecture. Multigene payloads that expressed robustly in permissive cell lines frequently failed in keratinocyte and fibroblast lineages despite correct integration, revealing regulatory constraints imposed by differentiated human cells. Promoter interference, inefficient ribosomal skipping, and cell-intrinsic signaling thresholds created silent bottlenecks - phenotypes in which therapeutic genes were present but biologically inactive. These findings shift the central challenge from delivery to regulation: multigene therapeutic constructs behave as interacting gene systems rather than independent expression units, and their functional efficacy is strictly governed by vector architecture and delivery modality (cellular context?).

### 3.1. Delivery Barriers and Site-Specific Integration

Our benchmarking data confirm that prime editing-directed integrase systems (eePASSIGE) offer a robust alternative to traditional homology-directed repair (HDR) for installing multi-kilobase payloads into safe harbors like *AAVS1* [25]. However, the translation of these tools from HEK293T models to therapeutically relevant skin cells presented a significant delivery barrier. While lipid-mediated transfection (Lipofectamine 3000) was efficient in HEK cells, it yielded negligible results in HaCaT and HDFn-hTERT lines. We overcame this by utilizing Nucleofection, which enabled nuclear uptake of the large prime-editing components. To mitigate the substantial cytotoxicity associated with this physical disruption, supplementation with the ROCK inhibitor Y-27632 was essential to suppress anoikis and support post-electroporation recovery [26]. This establishes a clear technical protocol: high-efficiency editing in skin progenitors requires aggressive nuclear delivery coupled with anti-apoptotic pharmacological support.

### 3.2. The “Architecture Effect”: IRES-Mediated Polycistronic Strategies

A pervasive challenge in synthetic biology is the “stacking” of multiple transcriptional units. Our results vividly illustrate transcriptional interference (TI), where a strong downstream promoter (CMV) occludes the activity of an upstream unit (EF1*α*) [21]. Furthermore, we observed that T2A ribosomal skipping peptides, while functional in HEK cells, failed in skin lineages, likely due to specific ribosomal pausing profiles or degradation of fusion products [18]. In stark contrast, the EMCV-IRES architecture proved robust across all cell types. Based on this, we propose that the optimal design for next-generation skin substitutes is a single-promoter polycistronic architecture or using large insulator sequence when using multiple TU design. This streamlined configuration leverages the proven efficacy of IRES elements in skin lineages to drive multiple genes from a single potent transcriptional start site. By abandoning multi-promoter stacks in favor of IRES-linked chains, future constructs can ensure stoichiometric co-expression of the metabolic checkpoint, the inhibitory ligand, and the safety switch without the silencing artifacts of promoter interference.

### 3.3. Safety Switches and Cell-Intrinsic Resistance

The inclusion of an inducible safety switch is a prerequisite for clinical translation. Our observation that full-length iCasp9 failed to induce apoptosis in HEK293T cells - despite robust mRNA expression - highlights the need for molecular refinement. The retention of the N-terminal CARD domain can sterically hinder drug-induced dimerization [22], while HEK-intrinsic Bcl-2 homologs (E1B-19K) buffer against mitochondrial depolarization [23, 24]. For clinical applications in primary keratinocytes, we recommend utilizing ΔCARD-iCasp9 variants to lower the activation threshold and ensure rapid clearance.

### 3.4. Context-Dependent Immunomodulation

Finally, our functional assays reveal the distinct nature of the IDO1 and PD-L1 pathways. IDO1 functions as a constitutive “metabolic sink,” though we confirm that high levels of IL-2 can override this starvation checkpoint [27]. Conversely, PD-L1 efficacy is context-dependent, requiring the expression of PD-1 on activated T-cells. In the clinical context of a severe burn, where infiltrating T-cells are naturally PD-1^*high*^ [10], we anticipate that PD-L1 will exert a dominant immunosuppressive effect. Therefore, the ideal “universal” skin graft must combine constitutive metabolic control (IDO1) with high-affinity checkpoint ligands (PD-L1), delivered via the optimized polycistronic architectures defined in this study.

Together, these observations define a general principle for engineered cell therapies: biological function emerges from the interaction between synthetic circuits and endogenous regulatory thresholds at transcriptional, translational, and signaling levels. Gene order establishes expression hierarchy, translational mechanisms determine protein availability, and cellular signaling state governs functional penetrance. In this framework, metabolic regulators such as IDO1 act as baseline gates, whereas receptor-mediated checkpoints such as PD-L1 operate as conditional amplifiers whose activity depends on immune activation context. The goal of engineering immune-evasive grafts is therefore not merely to introduce therapeutic genes, but to align circuit architecture with the regulatory logic of the host cell. These design constraints extend beyond skin substitutes and are broadly applicable to the construction of stable, multi-gene human cell therapies.

## 4. Materials and Methods

### 4.1. Reagents and Biological Materials

Cell culture reagents, including Dulbecco’s Modified Eagle Medium (DMEM), Dulbecco’s Phosphate-Buffered Saline (DPBS), TrypLE™ Express Enzyme, the ROCK inhibitor Y-27632, and Lipofectamine™ 3000, were obtained from Fisher Scientific. Fibroblast Growth Medium (FGM) and associated supplements were purchased from Cedarlane. Fetal Bovine Serum (FBS) was sourced from Corning and ATCC.

For molecular cloning and gene expression analysis, Q5® High-Fidelity DNA Polymerase, cDNA synthesis kits, SYBR® Green qPCR master mixes, and Gibson Assembly® HiFi Master Mix were purchased from New England Biolabs (NEB). Custom DNA oligonucleotides—including primers for cloning, qPCR, junctional PCR, and Site-Directed Mutagenesis (SDM)—were synthesized by Integrated DNA Technologies (IDT). Plasmids encoding PASTE and PASSIGE components, as well as donor backbone vectors, were obtained from Addgene.

Nucleic acid extraction was performed using Mini prep kits (Luna Nanotech) and Endotoxin-Free Midi/Maxi prep kits (Qiagen) for transfection-grade DNA. Genomic DNA and total RNA were isolated using the DNeasy® Blood & Tissue Kit and RNeasy® Mini Kit, respectively (Qiagen). Functional assays utilized the Indoleamine 2,3-Dioxygenase 1 (IDO1) Activity Assay Kit (Abcam). Fluorophore-conjugated antibodies, including PE-Cy7 anti-human CD274 (PD-L1), were purchased from BioLegend or BD Biosciences. A complete list of primer sequences, genome targeting sites, and plasmid maps is provided in the Supplementary Data (Supplementary Tables S1–S3).

### 4.2. Plasmid Construction and Minicircle Generation

Parental donor plasmids containing the *attP*-Bxb1 integration site were constructed via Gibson Assembly. Twin-PE gRNA plasmids targeting the *AAVS1* safe harbor locus were designed and cloned into the pU6-tevopreq1-GG-acceptor backbone (Addgene #174038).

To generate the evolved serine integrase expression vector (pCMV-eeBxb1), the wild-type *Bxb1* coding sequence was first amplified from the PASTE v3 template (pDY1052; Addgene #179105) and cloned into a standard pCMV expression vector. Subsequently, site-directed mutagenesis was performed to introduce the high-efficiency mutations V74A, E229K, and V375I into the *Bxb1* sequence.

For minicircle DNA production, donor cassettes were cloned into the pDY0181 backbone (Cargo EGFP with *attP*-Bxb1; Addgene #179115) and transformed into ZYCY10P3S2T *E. coli* cells (System Biosciences). Following arabinose induction to trigger PhiC31-mediated recombination and excision of the bacterial backbone, the resulting minicircle DNA was isolated using Qiagen Endotoxin-Free Maxi Prep kits.

### 4.3. Cell Culture

All cell lines were maintained at 37°C in a humidified 5% CO_2_ and appropriate humidity atmosphere without antibiotics. HEK293T cells were cultured in DMEM supplemented with 10% FBS. HaCaT keratinocytes (Applied Biological Materials (abm) Inc.) were maintained in DMEM supplemented with 10% FBS. Immortalized human neonatal dermal fibroblasts (HDFn-hTERT) (abm Inc.) were expanded in Fibroblast Growth Medium (FGM) supplemented with 10% FBS.

### 4.4. Genomic Integration Setup

Site-specific integration was achieved using the eePASSIGE platform, which co-delivers the prime editing machinery (pCMV-PEmax; Addgene #174820), a custom-evolved serine integrase (pCMV-eeBxb1), and dual pegRNAs (pU6-pegRNA1-AAVS1-attB, pU6-pegRNA2-AAVS1-attB) to install a genomic attB landing pad, alongside an attP-containing minicircle donor. Multi-kilobase donor cassettes included: Construct 637 (EF1α→IDO1-T2A-GFP), Construct 702 (EF1α→PD-L1–IRES–GFP), Construct 706 (EF1α→IDO1-BGH; CMV→PD-L1-SV40pA), Construct 715 (EF1α→IDO1-BGH; CMV→GFP-SV40pA), Construct 717 (EF1α→IDO1-BGH; CMV→PD-L1–IRES–DmrB-iCasp9-SV40pA).

Recombination Strategy: Donor minicircles bearing an attP site were co-delivered with the evolved serine integrase (eePASSIGE configuration) to catalyze attB×attP recombination. For benchmarking, PASTE and eePASTE methods were tested under matched delivery conditions using the PASTE v3 vector (pDY1052; Addgene #179105) or a custom vector encoding the triple-mutant Bxb1 integrase. Correct insertion was verified by junction-spanning PCR (attL/attR) and Sanger sequencing. pegRNA sequences and donor construct details are listed in Supplementary Tables S1–S3.

#### 4.4.1. Transfection and Nucleofection

HEK293T: Cells were transfected using Lipofectamine™ 3000 according to the manufacturer’s protocol.

Primary-Like Cells: HaCaT and HDFn cells were engineered using the 4D-Nucleofector™ System (Lonza). HaCaT cells were nucleofected using the P3 Primary Cell 4D-Nucleofector™ X Kit, and HDFn cells using the P2 Primary Cell 4D-Nucleofector™ X Kit. Immediately post-nucleofection, cells were resuspended in medium supplemented with the ROCK inhibitor Y-27632 (10 μM) for 24 hours to enhance recovery and survival.

### 4.5. Flow Cytometry and Cell Sorting

Engineered populations were enriched for transgene integration using Fluorescence-Activated Cell Sorting (FACS). Cells were harvested, washed in FACS buffer, and sorted based on GFP fluorescence or surface PD-L1 expression (stained with PE-Cy7 anti-human CD274). Sorting was performed on a SONY SH800 or BD FACSMelody™ cell sorter. Three consecutive rounds of bulk sorting were conducted to generate stable, high-purity polyclonal lines. Subsequently, single-cell sorting was performed for HEK293T and HaCaT engineered lines to isolate clonal populations.

### 4.6. PBMC Source, Activation, Labeling, and Co-Culture

Cryopreserved human PBMCs (single lot; STEMCELL Technologies) were thawed per the manufacturer’s instructions and rested for 24 hours. T cells were activated with anti-CD3/CD28 magnetic beads for 48 hours in medium containing recombinant human IL-2 (100 U/mL or 1000 U/mL), then rested for 24 hours. Immediately prior to co-culture, PBMCs were labeled with CellTrace™ Yellow (CTY) to track proliferation. Engineered or control HEK293T monolayers (grown to 70–90% confluency overnight) were co-cultured with labeled PBMCs for 3 days in 24-well plates at a ratio of 1:5 (3.0 x 10^5^ HEK to 1.5 x 10^6^ PBMCs). Controls included PBMCs alone (CTY-stained and unstained) and PBMCs co-cultured with wild-type HEK cells. Exogenous IL-2 (20 U/mL or 200 U/mL) was maintained during the 3-day co-culture period.

For flow Cytometric Analysis, T-cell proliferation was quantified by CTY dye dilution within viable T-cell gates. Specific T-cell subsets were identified using the following panel: anti-CD3 V500 (Lineage), anti-CD4 V450 (Helper T cells), anti-CD8 PE-Cy7 (Cytotoxic T cells), anti-CD25 APC (Activation/Treg), and anti-CD127 APC-eFluor 780 (Treg exclusion). Compensation beads (UltraComp eBeads™) and fluorescence minus one (FMO) controls were used to set gates.

### 4.7. RNA Extraction, cDNA Synthesis, and RT-qPCR

Total RNA was extracted using the RNeasy® Mini Kit (Qiagen) and reverse transcribed to cDNA. qPCR was performed on an Applied Biosystems real-time PCR system using SYBR® Green chemistry, with technical triplicates per sample. Primer efficiencies were validated, and analyses adhered to MIQE guidelines.

Given the absence of detectable endogenous IDO1 and PD-L1 expression in parental HEK293T, HaCaT, and HDFn-hTERT cell lines, transcript abundance was quantified relative to stable housekeeping genes rather than as a fold-change over wild-type controls. Target gene cycle threshold (Ct) values were normalized to the geometric mean of two reference genes, GAPDH and B2M *C*_*t*,*ref*.*geo*_). Relative expression was calculated as a percentage of reference gene expression using the formula:

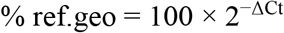

Non-template controls were confirmed negative; undetected targets were reported as “ND” (Not Detected).

### 4.8. Genomic DNA Extraction and Junction PCR

Genomic DNA was isolated using the DNeasy® Blood & Tissue Kit (Qiagen). Site-specific integration was assessed by 5′ and 3′ junction-spanning PCR to detect attL/attR recombination products and to exclude parental attP sites. Amplicons were verified by Sanger sequencing. Primer sequences and genomic coordinates are provided in Supplementary Tables S1–S3.

### 4.9. Functional Characterization

#### 4.9.1. IDO1 Enzymatic Activity

Kynurenine production was quantified using the IDO1 Activity Assay Kit (Abcam) according to the manufacturer’s instructions.

#### 4.9.2. Safety Switch Activation

Cells were treated with the dimerizer rimiducid (AP1903) at concentrations ranging from 1 nM to 1 μM. Cell viability was assessed 24 hours post-treatment via Annexin V/Propidium Iodide staining or metabolic viability assays (e.g., MTS/CellTiter-Glo).

### 4.10. Statistics and Reproducibility

Biological replicate numbers (n) and statistical tests are reported in the figure legends. Where multiple PBMC donors were used, donor identity was treated as a random factor in mixed-effects analyses. Data are presented as mean ± SEM or SD as indicated.

## Acknowledgments

This research was supported by the University of Manitoba Ignite Program. We gratefully acknowledge the Aleeza Gerstein Lab and the Rady Faculty of Health Sciences for their support with fluorescence-activated cell sorting.

## References

1. Shpichka, A., et al., Skin tissue regeneration for burn injury. Stem Cell Research Therapy, 2019. 10(1): p. 94.

2. Vig, K., et al., Advances in Skin Regeneration Using Tissue Engineering. Int J Mol Sci, 2017. 18(4).

3. Shevchenko, R.V., S.L. James, and S.E. James, A review of tissue-engineered skin bioconstructs available for skin reconstruction. Journal of Royal Society Interface, 2010. 7(43): p. 229–58.

4. Dixit, S., et al., Immunological challenges associated with artificial skin grafts: available solutions and stem cells in future design of synthetic skin. J Biol Eng, 2017. 11: p. 49.

5. Deuse, T., et al., Hypoimmunogenic derivatives of induced pluripotent stem cells evade immune rejection in fully immunocompetent allogeneic recipients. Nature Biotechnology, 2019. 37(3): p. 252–258.

6. Munn, D.H., et al., Prevention of allogeneic fetal rejection by tryptophan catabolism. Science, 1998. 281(5380): p. 1191–3.

7. Mellor, A.L. and D.H. Munn, IDO expression by dendritic cells: tolerance and tryptophan catabolism. Nat Rev Immunol, 2004. 4(10): p. 762–74.

8. Platten, M., et al., Tryptophan metabolism as a common therapeutic target in cancer, neurodegeneration and beyond. Nat Rev Drug Discov, 2019. 18(5): p. 379–401.

9. Pardoll, D.M., The blockade of immune checkpoints in cancer immunotherapy. Nat Rev Cancer, 2012. 12(4): p. 252–64.

10. Sharpe, A.H. and K.E. Pauken, The diverse functions of the PD1 inhibitory pathway. Nat Rev Immunol, 2018. 18(3): p. 153–167.

11. Hockemeyer, D., et al., Efficient targeting of expressed and silent genes in human ESCs and iPSCs using zinc-finger nucleases. Nat Biotechnol, 2009. 27(9): p. 851–7.

12. Anzalone, A.V., et al., Search-and-replace genome editing without double-strand breaks or donor DNA. Nature, 2019. 576(7785): p. 149–157.

13. Yarnall, M.T.N., et al., Drag-and-drop genome insertion of large sequences without double-strand DNA cleavage using CRISPR-directed integrases. Nature Biotechnology, 2023. 41(4): p. 500–512.

14. Straathof, K.C., et al., An inducible caspase 9 safety switch for T-cell therapy. Blood, 2005. 105(11): p. 4247–54.

15. Di Stasi, A., et al., Inducible apoptosis as a safety switch for adoptive cell therapy. N Engl J Med, 2011. 365(18): p. 1673–83.

16. Gossen, M. and H. Bujard, Tight control of gene expression in mammalian cells by tetracycline-responsive promoters. Proc Natl Acad Sci U S A, 1992. 89(12): p. 5547–51.

17. Das, A.T., L. Tenenbaum, and B. Berkhout, Tet-On Systems For Doxycycline-inducible Gene Expression. Curr Gene Ther, 2016. 16(3): p. 156–67.

18. Kim, J.H., et al., High cleavage efficiency of a 2A peptide derived from porcine teschovirus-1 in human cell lines, zebrafish and mice. PLoS One, 2011. 6(4): p. e18556.

19. Eszterhas, S.K., et al., Transcriptional interference by independently regulated genes occurs in any relative arrangement of the genes and is influenced by chromosomal integration position. Mol Cell Biol, 2002. 22(2): p. 469–79.

20. Gill, G. and M. Ptashne, Negative effect of the transcriptional activator GAL4. Nature, 1988. 334(6184): p. 721–4.

21. Shearwin, K.E., B.P. Callen, and J.B. Egan, Transcriptional interference--a crash course. Trends Genet, 2005. 21(6): p. 339–45.

22. Straathof, K.C., et al., An inducible caspase 9 safety switch for T-cell therapy. Blood, 2005. 105(11): p. 4247–54.

23. White, E., Regulation of the cell cycle and apoptosis by the oncogenes of adenovirus. Oncogene, 2001. 20(54): p. 7836–46.

24. Zhang, W., et al., Generation of apoptosis-resistant HEK293 cells with CRISPR/Cas mediated quadruple gene knockout for improved protein and virus production. Biotechnol Bioeng, 2017. 114(11): p. 2539–2549.

25. Pandey, S., et al., Efficient site-specific integration of large genes in mammalian cells via continuously evolved recombinases and prime editing. Nat Biomed Eng, 2025. 9(1): p. 22–39.

26. Watanabe, K., et al., A ROCK inhibitor permits survival of dissociated human embryonic stem cells. Nat Biotechnol, 2007. 25(6): p. 681–6.

27. Munn, D.H. and A.L. Mellor, Indoleamine 2,3 dioxygenase and metabolic control of immune responses. Trends Immunol, 2013. 34(3): p. 137–43.

